# Protein kinase J regulates Rv0642c expression and sensitivity to rifampicin of mycobacteria

**DOI:** 10.1101/2021.06.13.448225

**Authors:** Diwakar Kumar Singh, Samir Tiwari, Kumari Kripalata, Kishore K. Srivastava

## Abstract

Mycobacterium tuberculosis (M.tb) persists for long-duration inside the human host in both active and latent form by modulating the immune response. The mechanisms employed by M.tb to survive inside the host and evade the host immune system need to be explored in greater depth for the rational design of novel treatment strategies. The phosphorylation and methylation of biomolecules need to be addressed in mycobacteria because it has an important role in infection establishment and persistence. In the present study, we elaxborate on the role of PknJ in the slow growth of BCG and its association with mmaA4 protein during extracellular and intracellular growth. The *pknJ* knock-out (KO) BCG has been used to decode the functional significance in mycobacteria. The mmaA4 expression and methyltransferase activity is decreased in knocked-out BCG strain (*pknJ*^-/-^) during extracellular growth, while mmaA4 expression and methyltransferase activity is increased during intracellular growth of mycobacteria. The knocked-out BCG strain is highly sensitive to the rifampicin antibiotics during extracellular growth in compared to control. A significant association of *pknJ* and *mmaA4* was found in our studies during the growth and intracellular persistence of mycobacteria.

## Introduction

The mycobacteria encounter several types of stresses during ligand treatment and disease manifestation. They have evolved different defence mechanism depending on kind of stress and cell type / tissue type. Impaired host immunity has been assumed to be a measure causative factor in tuberculosis infection. The signalling pathway of Mycobacterium tuberculosis are controlled by 11 serine threonine protein kinase (STPK), one Ser/Thr phosphatase, one tyrosin kinase and two tyrosine phosphatase (Bach et al., 2009; Cole et al., 1998). The phosphorylations involved in mycobacteria and regulate the cell wall biosynthesis. The kinases and phosphatases are molecular switch for translating extracellular signals into cellular responses. The phosphorylation of protein is associated with the virulence factor, affects the mycobacterial physiology (Av-Gay and Everett, 2000). The several investigations have suggested, expression modulation of serine threonine protein kinase changes the pathological and physiological significance of mycobacteria. Like, cyclopropane synthase PcaA was phosphorylated by STPK and modulate the intracellular survival of mycobacteria. Further, pcaA mutant showed defect in mycolic acid modification in mycobacteria and unable to persists within host (Corrales et al., 2012; Glickman et al., 2000). Over-expression of PknA in *Mycobacterium bovis* BCG results in a cells forming an elongated, branched structure, whereas overexpression of PknB results in the formation of widened and bulging cells (Kang et al., 2005). Similarly, PknA and PknB phosphorylate several proteins including Wag31 (Rv3083) (Kang et al., 2005), a homolog of the cell shape / cell division protein DivIV which are essential for mycobacterial growth (Nguyen et al., 2007). The penicillin binding proteins of mycobacteria viz, PbpA (Rv0016c) is phosphorylated by PknB (Dasgupta et al., 2006). The phosphorylation of VirS by PknK controls biosynthesis of mycolic acid and important for maintaining the cell wall integrity and virulence in mycobacteria(Qu et al., 2021; Singh et al., 2005b).

Besides of serine threonine protein kinases, pathogenic mycobacteria can manipulate their mycolate profiles to adapt in different environment during the host infection and stress response. The selective expression and secretion of mycolic acids implies their requirement during the tuberculosis infection of the host (Verschoor et al., 2012). Mycolic acids are essential for intracellular survival of mycobacteria, resistant to broad-spectrum antimicrobials (Marrakchi et al., 2014; Yuan et al., 1998). The H37Rv intra cellular infection of macrophage leads to increased synthesis of ketomycolate (Yuan et al., 1998). The enhanced ketomycolate production (mmaA4 gene) leads to increased survival of mycobacteria during stress response, host infection and biofilm formation (Dubnau et al., 2000; Sambandan et al., 2013; Yuan et al., 1998). *Mycobacterium tuberculosis* has many SAM dependent methyltransferases (SAM–MTs) which participate in mycolic acid modification like; *mmaA*1-4, *cmaA*1-2, *pcaA* and *umaA* (Barkan et al., 2012). SAM-MTs are enzymes which modify the cyclopropanation and introduce methyl branches in mycolic acids. The STPK has strong association in modulation of mycolate synthesis during mycobacterial infection.

In the present study we have shown the regulatory role of *pknJ* in *mmaA4* expression which is important component of mycolic acid. We have confirmed, PknJ retard the growth of mycobacteria after knocking down its expression (Singh et al., 2014). Now we have extended our studies using knockout BCG strain (*pknJ*^-/-^) in MGIT 960 and macrophage cells. The knock-out strain has defect in mmaA4 expression in both intracellular and extracellular condition. The methyltransferase activity and rifampicin sensitivity is interrelated inside the bacterial cells and can be regulated through *pknJ* expression which modulated mmaA4 expression. The rifampicin antibiotics has association with mycolic acid (Sambandan et al., 2013; Yuan et al., 1998). Since, this study fills the lacuna of mmaA4 candidate gene in mycobacteria.

## Materials and methods

### Bacterial cultures and Cell line

Mycobacterial strains, *Mycobacterium bovis* BCG laboratory strain (ATCC#35734, TMC 1011 BCG Pasteur) and *M. tuberculosis* H37Rv (ATCC#25618) were procured through ATCC (USA). *Escherichia coli* (*E. coli*) strains DH5α and BL21 (DE3) were procured from NEB and were cultured in Luria–Bertani medium. The murine macrophage cell line J774A.1 was obtained from the American Type Culture Collection (ATCC), USA, and cultured in RPMI-1640 medium (2 mML-glutamine,10 mM HEPES, 1 mM sodium pyruvate, 4.5 g/L glucose and 1.5 g/L sodium bicarbonate), supplemented with 10 % heat-inactivated fetal calf serum (FCS) at 37 °C and 5 % CO2.

### Cloning, expression, and purification of MTB Rv0642c

The *mmaA4* (Rv0642c) gene was PCR-amplified from the genomic DNA of MTB H37Rv using gene-specific primers (Table 1). *Hind*III restriction sites were incorporated at the 5’end of forward and reverse primers. The PCR-amplified product was cloned and expressed in *E. coli* BL21 (DE3) cells by 0.2mM IPTG at 22°C for overnight. The expressed proteins were purified with Ni-NTA column and gel filtration chromatography [17]. The integrity of purified protein was checked on SDS PAGE and was found at 34.7 kDa. The *pknJ* was subcloned into modified episomal pMV261 vector, inserted *E.coli* ori by replacement of mycobacterial ori (Singh et al., 2015). This is used for mutant/knockout BCG development and *pknJ* was further sub cloned into pMV361 used for complementation of mutant BCG strain. The cloning and purification of *pknJ* was reported earlier (Singh et al., 2014).

### Generation of polyclonal antisera in rabbit and immunoblot assay

Polyclonal antisera were raised and purified by Protein G Sepharose for immunoblotting assay as described previously [17; 18]

### RNA isolation and quantitative RT-PCR (qRT-PCR)

The total RNA was extraction and qRT-PCR as described previously [17]

### In vitro kinase assay using MBP and MmaA4 as substrate

In vitro kinase assays were performed by ADP-glo kinase assay using 30ng of recombinant PknJ (rPknJ) with 25, 50 and 100ng of _r_mmaA4 protein at 24 °C for 1 h as described earlier (Singh et al., 2014).

### Infection of J774A.1 macrophage cells with recombinant BCG

The J774A.1 macrophage cells was cultured in RPMI-1640 medium supplemented with 10% heat inactivated fetal calf serum (FCS) at 37 °C and 5 % CO2, 5×x10^5^ cells/ml was seeded in tissue culture plates, incubated overnight and used for BCG infection as described previously (Singh et al., 2014).

### Methyltransferase activity in recombinant BCG

The BCG log and stationary phase cultures were used for methyltransferase activity by colorimeter. Wild-type BCG and mutant/knockout BCG were lysed harvested according to standards protocol. The soluble protein extracts was separated by centrifugation at 12,000×g for 30 min and filtered through a 0.8 μm. Twenty micrograms of total soluble proteins were used for enzymatic activity at different stages of mycobacteria. The enzymatic reaction was carried out according to manufacturer instruction of Methyltransferase Colorimetric Assay Kit (Cyman Chemical). The reactions were performed at 37 ^0^C to monitor the change in absorbance at 510nM using plate reader. The background reaction was calculated in absence of SAM as well as in absence of total protein in reaction mixture. The enzymatic reaction was monitored continuously up to 10 minutes and determined the rate of production of hydrogen peroxide by colorimeter at 510 nm.

### Bacterial culture and rifampicin (RIF) susceptibility by MGIT 960 and MABA

BCG, mutant BCG was cultured in Middlebrook 7H9 medium (Difco) supplemented with Albumin Dextrose Catalase (BD). Mid log phase culture was used to prepare the suspension of bacilli by repeated vortexing in Middlebrook 7H9 broth. The cultures were diluted 1:100 before the bacilli suspension and retested by the CFU counting on solid plates as described (Singh et al., 2014; Singh et al., 2015). The three strains growth was monitored up to 12 days in BACTEC MGIT960 system. The data was recorded every day and stored for further analysis. Further 2ug RIF were tested for growth of BCG and mutant BCG under the similar condition in BACTEC MGIT960 system.

To revalidate the 1st line drug susceptibility (RIF) of mutant BCG in MGIT960 system. We elaborated our study for the MABA assay. The antimicrobial testing was performed in 96-well microplates at 37°C. The different concentration of RIF (0.125 to 16 ug) was tested in the bacilli suspension and monitored the growth of bacteria. The alamar blue solutions were used to test the colour change in test and control sample at 12 and 24h. The fluorescence was further measured in fluorometer with excitation at 530nm and emission at 590nm.

### Protein isolation and labelling with Cy dyes in 2D gel electrophoresis

The protein isolation and labelling as described previously (Singh et al., 2014).

## Result

### PknJ phosphorylates mmaA4 and modulates expression and intracellular survival of mycobacteria

To identify the mmaA4 phosphorylation by PknJ, both proteins were heterogeneously expressed and purified as shown in (Fig. 1A). The kinase assay was done using the purified recombinant proteins. The kinetically active PknJ protein phosphorylates mmaA4 recombinant protein in cell free system (Fig. 1.B and 1.C). The negative control was taken as a kinase reaction in absence of specific substrate and luminescence was measured at the end reaction. The mutant BCG strain (*pknJ*^-/-^) was developed using strategies (Fig. 1D). PknJ expression was determined in BCG and knockout BCG using polyclonal PknJ hyper immune sera (Fig. 1E). The mmaA4 encoding gene is present in both pathogenic and non-pathogenic strains of mycobacteria. The deficient expression of *mmaA4* gene was observed in knockout BCG while restored after *pknJ* complementation (Fig. 1F). The in-vitro culture of both strains was used for 2D in gel analysis (Fig. 1H). The 3D image of same spots was identified in 2D-DIGE analysis and shown adjacent to gel in same figure (Fig. 1H). The selected spot was reconfirmed by LC-MS and identified peptides shown as an underlined in sequence (Fig. 1G). The BCG and knockout BCG (*pknJ*^-/-^) was cultured in sautons medium, used to infect the J774A.1 cell line. The intracellular survival of mycobacteria was monitored up to 96h. The result showed that mycobacteria containing *pknJ* increased the survival in comparison to others (Fig. 1I). The both strains showed the significant difference in survival during 24h to 96h of J774A.1 infection.

**Fig. 1.**
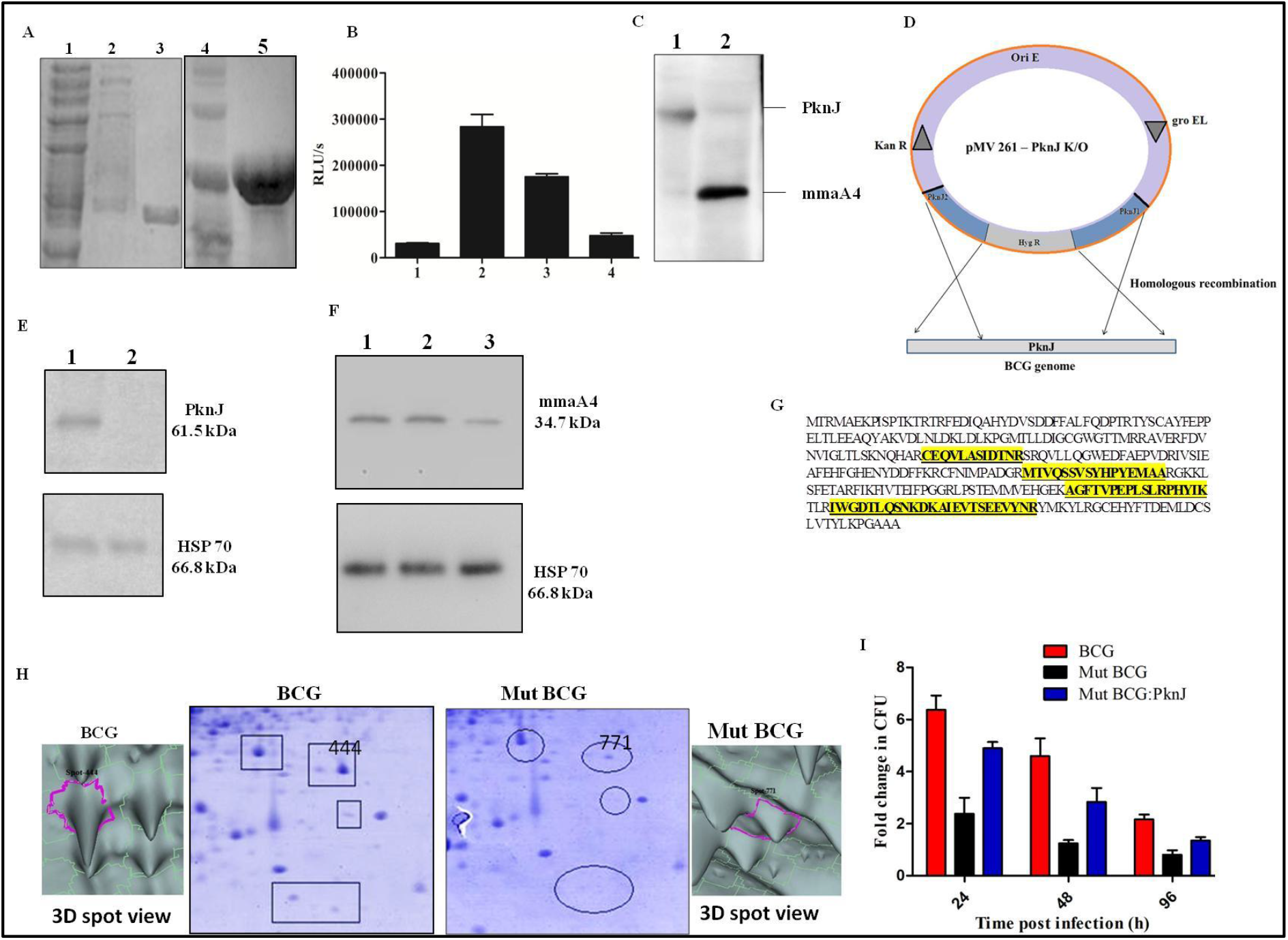
PknJ down regulates the expression of mmaA4 in intracellular survival: A. Purification of mmaA4 and PknJ proteins with affinity and gel filtration chromatography. Lane1-flow-through of cell lysates, lane2- wash of flow-through, lane3- purification of recombinant mmaA4 protein (34.7 kDa), Lane4- flow-through of cell lysates, Lane5- purification of PknJ (61.5kDa) : B. PknJ kinase assay using MBP and different concentrations of rmmaA4 as a substrate. Lane1- MBP as a substrate, lane 2- 100 ng rmmaA4 protein, lane 3- 50 ng rmmaA4 protein and lane 4- 25 ng rmmaA4 protein: C. phospho serine/threonine antibody was used in immunobloting to detect the phosphorylation of kinase reaction. Lane 1- phosphorylation MBP of PknJ, lane 2- upper band is autophosphorylation of PknJ and lower is phosphorylation of mmaA4: D. Strategy and diagrammatical representation for generation of knockout BCG (*pknJ*^-/-^) : E. BCG and knockout BCG cell lysate proteins (10ug) were immunoblotted with for α-PknJ (hyperimmune sera). Lane 1- BCG and Lane 2- knockout BCG: F. BCG and knockout BCG cell lysate (10ug) were used for immunoblotting with mmaA4 hyperimmune sera. The mmaA4 expression is same in BCG, Lane1, complemented BCG, Lane 2 while diminished in knockout/mutant BCG, Lane 3: G. the 2D gel spot was identify/ sequenced by MALDI and identified peptide is underlined: H. The BCG and knockout BCG protein analysed by in gel analysis and mmaA4 spot is marked by arbitratory spot number (444 in BCG and 771in mutant BCG) and encircled. The 3D spot view was identified by instrumental software and shown adjacent to the image: I. Both BCG strains were used for survival assay in J774A.1 cells. The infected cell were lysed and plated on MB7H10 plates containing 10%OADC and 25 μg/ml kanamycin according to given time frame. The CFU counts were done at different time interval and determined the fold change in CFU (24, 48, and 96h) from three independent experiments in three replicates.

### Intracellular methyltransferase activity in response to PknJ and Rv0642c expression

To investigate whether methyltransferase activity is altered in intracellular BCG and knockout BCG because it has important role in mycolate synthesis and virulence. The PknJ has role in alteration of morphology according to our previous report by electron microscopy (Singh et al., 2014). The methyltransferase activity was optimised using the positive control and BCG cell lysate (Sup Fig. 1 & 2). We measured methyltransferase activity in logarithmic cultures of both strains and found knockout has lower enzymatic activity under the similar experimental condition (Fig. 2A). The less enzymatic activity is due to poor expression of mmaA4 in mutant/ knockout strain (Fig. 1F). In order to elaborate the study and checked enzymatic activity during the stress response of cell. We found dividing mycobacteria have high enzymatic activity because cell wall associated component requires more to compensate the growth condition (Fig. 2B). We have analysed the transcript of mmaA4 by real time pcr using gene specific primer given in table 2 and found high level of transcript during the stationary phase (Fig. 2C). It is not surprising because mmaA4 gene is responsible for synthesis of methoxy and keto mycolate and it protect the mycobacteria against various stress condition. The mmaA4 is major components required for survival of intracellular mycobacteria in macrophage cells. We have tested this hypothesis by immunobloting and found increased level of mmaA4 expression in mutant BCG compared to BCG during 48h of infection (Fig 2E). However, methyltransferase activity was high in intracellular mutant BCG in compare to BCG at 24, 48 and 72h (Fig 2D). The mycobacteria try to survive in compromised condition within the host by increasing the methyltransferase activity because methoxy mycolate and ketomycolate is essential component for its survival.

**Fig. 2.**
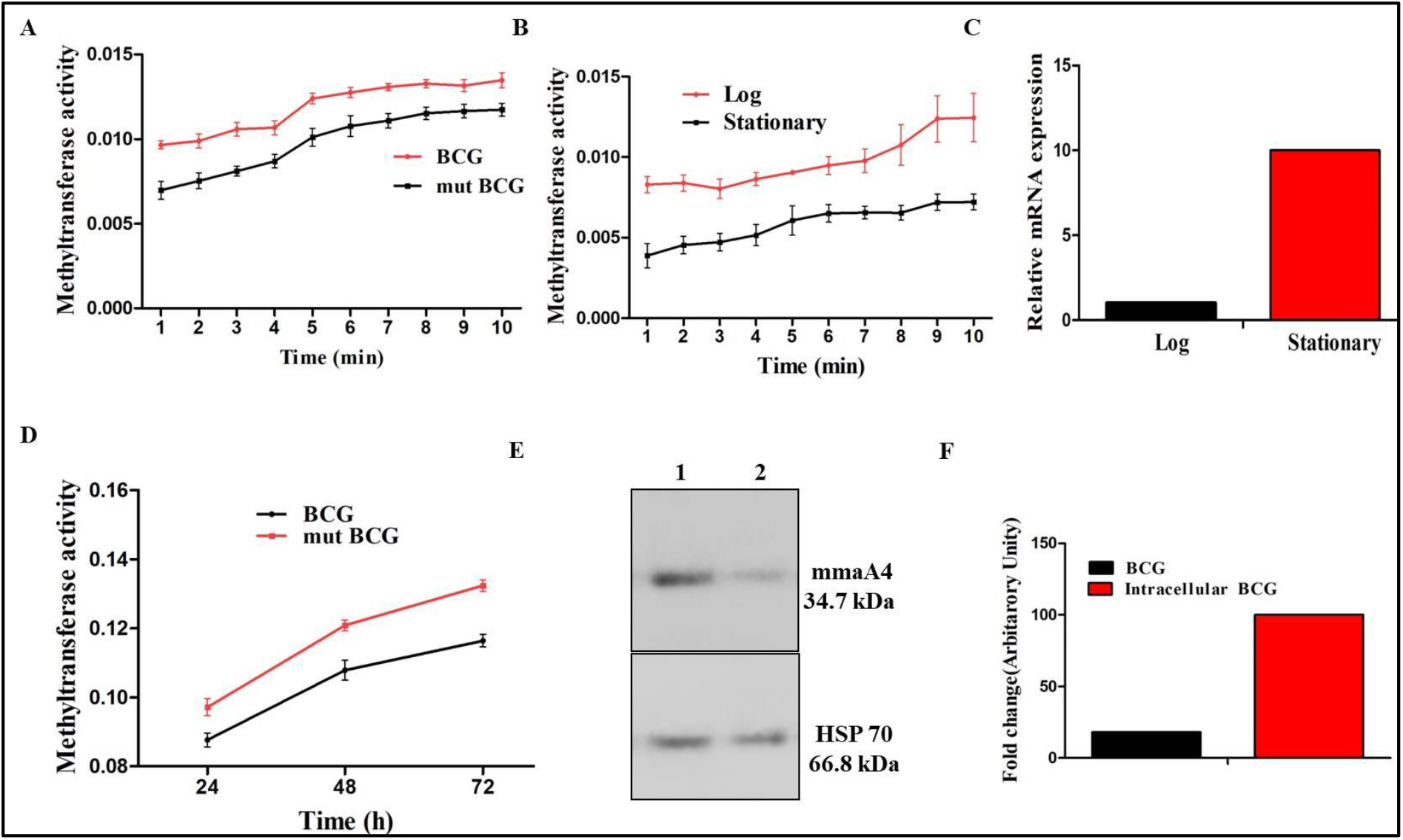
Methyltransferase activity in BCG. A. The protein was harvested, quantified from log-phase of BCG, mutant BCG and used equally for enzymatic activity (μmole/min): B. The log-phase culture were allowed to grow stationary phase and equal amount of protein was used for enzymatic activity: C. The transcript level of mmaA4 at log and stationary phases of BCG: D. The BCG and mutant BCG was recovered from infected J774A.1 cells at different time (24, 48 and 72h) and equal amount of proteins ware used for enzymatic activity. The enzymatic activity was monitored continuously up to 10 minutes. The single time point indicated above is average values of 10 minutes: E. the intracellular mycobacteria, 48h of infection were harvested and equal concentration of proteins was used for mmaA4 and hsp70 immunoblotting. Lane 1- Intracellular mutant BCG cell lysate and Lane 2- BCG cell lysate. The same blot was used for hsp70 probing as a loading control: F. The intensity of band was quantified by image J software. The mmaA4 band intensity were normalised by hsp 70 band intensity before fold change calculation.

### Antibiotic susceptibility test of mycobacterial culture

Mycobacteria have unique fatty acid synthesis by Fas1 and Fas II pathway. The complex mycolic acid structure of mycobacteria helps in antibiotics resistance during the treatment. The first line tuberculosis drug rifampicin (RIF) acts on the mycolic acid synthesis pathway and inhibits growth. Hence it was required to test our constructed BCG for the same. Initially, we checked the growth response of mycobacteria (BCG and mutant BCG) by CFU counting, in MGIT960 system (Fig. 3A). The mutant BCG showed higher growth in the cell-free system and restored the growth maximally after complementation with the same *pknJ* ORF (Fig. 3A).

**Fig. 3.**
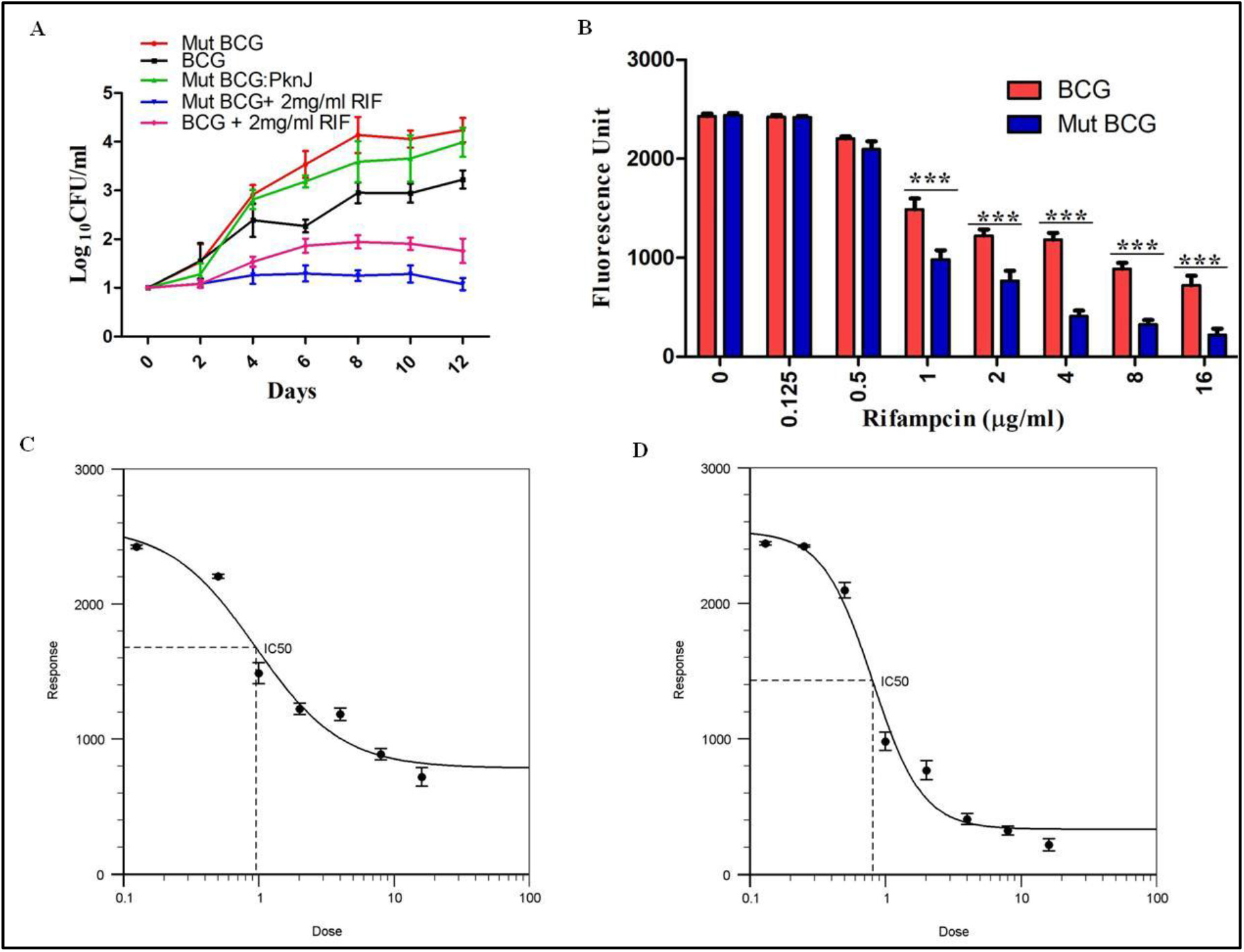
Growth kinetics of knockout/Mut BCG : A. Growth of BCG, Mut BCG (*pknJ^-/-^*) and complimented with *pknJ* (Mut BCG: *pknJ*) were monitored by CFU counting. The experiment was repeated thrice in triplicates. The statistical significance was checked by two-way ANOVA. Mutant BCG and BCG (p<0.0001, ***) during 6, 8, 10 and 12 days. The mutant BCG is more sensitive to rifampicin (2mg/ml) in compare to BCG strain. The both strains shows significant difference at 6, 8, 10 and 12 days (p>0.0001, **): B. The different concentration of rifampicin was used to monitor the growth kinetics of BCG and mutant BCG by MABA assay. More than 1ug/ml of rifampicin shows significant difference in growth between BCG and Mut BCG and all are statistically significant (p<0.0001, ***): C. IC50 of rifampicin against the BCG and mutant BCG was calculated by AAT Bioquest software and is shown above. IC50 of BCG= 0.927, IC50 of mut BCG= 0.803.

To test whether both s’ growth was affected by RIF treatment, we critically monitored it by CFU counting and MGIT960 system. The 2ug/ml RIF showed mutant s are more sensitive to BCG (Fig. 3A). The same culture was treated with different concentration of RIF and used for MABA test. The significant defect in growth was observed in 1 to 16ug/ml of antibiotic (Fig. 3B). IC50 was calculated by AAT Bioquest software of both strains (Fig. 3C, 3D)

## Discussion

We highlight the association of *Mycobacterium tuberculosis* STPK, PknJ and kinase substrate mmaA4 during growth of mycobacteria. The mmaA4 gene is responsible for synthesis of oxygenated component of mycolate and maintains the proper cell wall function during the in vivo growth of MTB (Yuan et al., 1998). The STPK of mycobacteria has been well documented in phosphorylation of protein involved in virulence, growth, cell division viz, cyclopropane synthase PcaA phosphorylation leads to intracellular survival (Corrales et al., 2012; Glickman et al., 2000), VirS phosphorylation control biosynthesis of mycolic acid (Singh et al., 2005a) and many of other key enzymes involved in mycolic acid synthesis pathway are substrates of mycobacterial STPK (Molle et al., 2006).

The importance of PknJ in growth and survival was shown in our earlier report, using BCG and *Mycobacterium smegmatis* (MS) as a surrogate strain (Singh et al., 2014). Now, we extended our studies and rectified the limitation of earlier report viz, gene knockdown. The mutant BCG (*pknJ*-/-) was developed through recombination process and used in this investgatation. We used BCG and mutant BCG as a plaktonic culture to revalidate the growth response of PknJ in mycobacteria. The extracellular growths of both strains were monitored by CFU counting as well as by MGIT960 system. We observed the mutant BCG has grown faster than wild type during extracellular culture and it restored the slow growth after complementation with BCG- *pknJ* (Fig. 3A). We are not surprising because it validates our previous observation in *pknJ* knockdown studies [16]. There are several reports, which have shown the importance of mmaA4 in intracellular survival of mycobacteria while its association with STPK is a still mater of investigation (Dubnau et al., 2000; Sambandan et al., 2013; Yuan et al., 1998). The mmaA4 phosphorylation by PknJ has been shown by Phospho-chip analysis of peptide phosphorylation by rPknJ (Jang et al., 2010). The Catechol O-methyltransferase (COMT) is transferred the methyl group onto hydroxyl groups of a catechol and enhance the biological significance in mycobacteria (Lee et al., 2019). The mmaA4 gene is present in pathogenic mycobacteria and their homologs are in non-pathogenic strains according to their genome data base. We have studied rmmaA4 phosphorylation by rPknJ in kinase reaction and specificity of phosphorylation was confirmed by the linearity in kinase activity when using different doses of the substrates (Fig. 1B). It was further reconfirmed by immunoblotting by using phospho serine/threonine antibody (Fig. 1C). The PknJ used mmaA4 as a substrate in our in-vitro analysis which is the initial sign of its in vitro association. The mycobacterial gene associations inside the cell have been studied using the mutant strain as a model during intracellular and extracellular growth (Fig. 1D). The mutant was confirmed by PCR and immunobloting (Fig. 1E). We found strong regulatory association between the genes. Hence, the mmaA4 expression in BCG and mutant BCG (*pknJ*^-/-^) was monitored by immunoblotting and 2D analysis and found decrease mmaA4 induction in mutant BCG strains (Fig. 1F, 1H). This mmaA4 regulatory defect arises due to *pknJ* deficiencies in mycobacteria. The mmaA4 induced the methoxy and ketomycolate synthesis which are essential for survival of mycobacteria during the infection and physiological stress condition (Dubnau et al., 2000; Sambandan et al., 2013; Yuan et al., 1998). The mmaA4 is an essential cell wall associated component required for pellicle growth, drug resistance and as a vaccine molecule (Sambandan et al., 2013). The methylation is involved in virulence and influences the mechanism of *M.tb.* pathogenesis (Grover et al., 2019). We checked the mmaA4 expression defects impacting upon the enzyme activity and observed SAM-MTs activity was inhibited in log phase culture of mutant BCG compare to wild type (Fig. 2A). This is obvious that the mmaA4 expression defect in mutant strains may lead to modulate enzymatic activity (Fig. 1F, 2A). We have elaborated the study and monitored SAM-MTs activity in log and stationary phage of mycobacteria and found high during the log phase culture (Fig. 2B). The bacteria are actively dividing in the log phase so may have a high demand of cell wall associated components. The posttranslational modification along with activation other gene may be the reason for decrease of enzymatic activities during the stress response. However, the expression of mmaA4 gene was high during stationary phage in qRT PCR analysis (Fig. 2C). The high expression of the mmaA4 gene may have a protective role in mycobacteria against the different stress in stationary phase viz, secretion of cellular metabolite and deficiency of oxygen etc.

In order to understand the survival defect of BCG and mutant BCG during the infection. We have used J774A, macrophage cell line as a host for infection studies and found less survival of mutant BCG (Fig. 1I). It was interesting to analyse the SAM-MTs activity of intracellular mycobacteria and found high in mutant BCG strain compared to BCG (Fig. 2D). However, the temporal expression of mmaA4 was also high in mutant BCG (Fig. 2E). The high mmaA4 expression and its enzymatic activity of intracellular mutant BCG (pknJ^-/-^) entails significant role of mmaA4 in the protection. The high expression and enzymatic activities may be due to deficiency of its kinase partner during the compromised growth of mycobacteria inside the host. The *pknJ* has great influence over the expression of mmaA4 during extracellular culture of BCG. We tested the potential effect of mmaA4 expression inside the cells through RIF sensitive assay. The RIF is a first line anti tubercular drug and inhibitor cell wall biosynthesis. Therefore easily define the biological significance of gene association in mycobacteria and found mutant is more sensitive to rifampicin antibiotics compare to wild type (Fig. 3A). The IC50 of mutant BCG is 0.803 and IC50 of BCG 0,927 was detected in MABA assay. The drug sensitivity/inhibitory effect suggesting an effect of *pknJ* on synthesis of oxygenated mycolate in *Mycobacterium tuberculosis*.

## Acknowledgements

We are thankful to the Director of CSIR-CDRI for access to the Sophisticated Analytical Instrumentation Facilities. The work has been supported by CSIR UNDO and CSIR-SPLENDED network projects. We thank Dr. J. K. Saxena for allowing us to use his facility for biochemical assay.

## Appendix A

Supplementary data

