## Supplementary for "Protein kinase J regulates Rv0642c expression and sensitivity to rifampicin of mycobacteria"

| Table 1. Primer sequence used in study | | |
|---|---|---|
| **A. Primer for cloning** | | |
| **Primer Name** | | **Primer Sequence (5'-3')** |
| PknJ-MTB | Forward | CCCAAGCTTGTGGCCCACGAGTTGAGTG |
| | Reverse | CCCAAGCTTTCAGCCGGGTATCTTGGCG |
| MmaA4-MTB | Forward | CCCAAGCTTATGACGAGAATGGCCGAGAAA |
| | Reverse | CCCAAGCTTTTAGGCCGCGGCACCCGG |
| **B. Primers for Real Time PCR** | | |
| MmaA4-MTB | Forward | ATGGCCGAGAAACCGATTAG |
| | Reverse | GGTCGACCTTGGCGTATTG |
| 16s rRNA | Forward | TGCAAGTCGAACGGAAAGGTCTCT |
| | Reverse | AGTTTCCCAGGCTTATCCCGAAGT |

Table 2. Up/down regulated spots detected by DIGE labelling.

| Position Number | Upregulated Proteins (Fold Change) | Downregulated Proteins (Fold Change) | Expected pI | Expected Mol.Wt (Kd) |
|---|---|---|---|---|
| 399 | 2.03 | | 4.2 | 68.3 |
| 404 | | 2.9 | 6.02 | 68.3 |
| 444 | | 10.67 | 6.68 | 62.6 |
| 450 | | 3.37 | 5.38 | 61.7 |
| 549 | | 3.49 | 6.16 | 51.08 |
| 557 | | 10.71 | 5.96 | 50.19 |
| 562 | | 5.86 | 5.76 | 50.3 |
| 574 | | 4.35 | 6.1 | 49.43 |
| 767 | | 6.74 | 6.09 | 33.5 |
| 771 | | 2.1 | 5.63 | 34.02 |
| 785 | | 2.96 | 5.87 | 34.67 |

Supplementary Fig.1: Methyltrasferase assay was optimized using a positive control in a reaction.

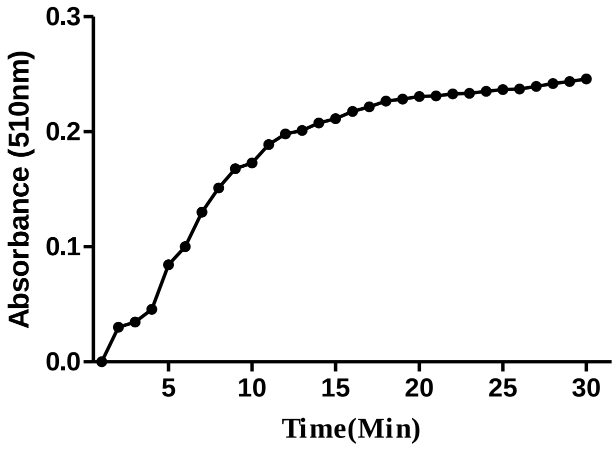

Supplementary Fig 2. Methyltrasferase assay was performed using the BCG cell lysate in presence of other components in a reaction and observed significantly high. Signal background 1 is in absence of protein and Signal background 2 is in absence of SAM.

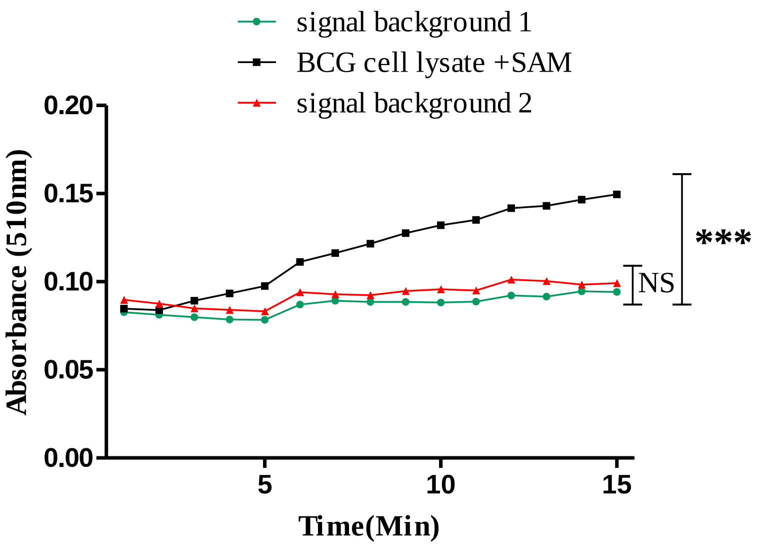
